# Joint sequencing of human and pathogen genomes reveals the genetics of pneumococcal meningitis

**DOI:** 10.1101/386078

**Authors:** John A. Lees, Bart Ferwerda, Philip H. C. Kremer, Nicole E. Wheeler, Mercedes Valls Serón, Nicholas J. Croucher, Rebecca A. Gladstone, Hester J. Bootsma, Nynke Rots, Alienke J. Wijmega-Monsuur, Elisabeth A. M. Sanders, Krzysztof Trzciński, Anne L. Wyllie, Aeilko H. Zwinderman, Leonard H. van den Berg, Wouter van Rheenen, Jan H. Veldink, Zitta B. Harboe, Lene F. Lundbo, Lisette C. P. G. M. de Groot, Natasja M. van Schoor, Nathalie van der Velde, Lars H. Ängquist, Thorkild I.A. Sørensen, Ellen A. Nohr, Alexander J. Mentzer, Tara C. Mills, Julian C. Knight, Mignon du Plessis, Susan Nzenze, Jeffrey N. Weiser, Julian Parkhill, Shabir Madhi, Thomas Benfield, Anne von Gottberg, Arie van der Ende, Matthijs C. Brouwer, Jeffrey C. Barrett, Stephen D. Bentley, Diederik van de Beek

**Affiliations:** Department of Microbiology, New York University School of Medicine, New York, United States of America; Infection Genomics, Wellcome Sanger Institute, Cambridge, United Kingdom; Department of Neurology, Amsterdam Academic Medical Centre, University of Amsterdam, Amsterdam, Amsterdam Neuroscience, The Netherlands; Department of Infectious Disease Epidemiology, Imperial College London, London, United Kingdom; Centre for Infectious Disease Control, National Institute for Public Health and the Environment, Bilthoven, The Netherlands; Department of Pediatric Immunology and Infectious Diseases, Wilhemina’s Children Hospital, University Medical Centre Utrecht, Utrecht, The Netherlands; Department of Biostatistics, Academic Medical Center, Amsterdam, The Netherlands; Department of Neurology, Brain Center Rudolf Magnus, University Medical Center Utrecht, Utrecht, The Netherlands; Department of Microbiological Surveillance and Research, Statens Serum Institut, Copenhagen, Denmark; Department of Infectious Diseases, Hvidovre Hospital, University of Copenhagen, Hvidovre, Denmark; Department of Human Nutrition, Wageningen University, PO Box 17 6700 AA Wageningen, The Netherlands; Amsterdam Public Health Research Institute, Department of Epidemiology and Biostatistics, VU University Medical Center, Van der Boechorststraat 7, 1081 BT Amsterdam, the Netherlands; Amsterdam Public Health Research Institute, Department of Internal Medicine, Geriatrics, AMC, Meibergdreef 9, 1105 AZ Amsterdam, The Netherlands; Department of Internal Medicine, Erasmus MC, University Medical Centre Rotterdam, P.O. Box 2040, 3000 CA Rotterdam, The Netherlands; Center for Clinical Research and Disease Prevention, Bispebjerg and Frederiksberg Hospitals, The Capital Region, Copenhagen, Denmark; The Novo Nordisk Foundation Center for Basic Metabolic Research, Section of Metabolic Genetics, Denmark; The Department of Public Health, Section of Epidemiology, Faculty of Health and Medical Sciences, University of Copenhagen, Copenhagen, Denmark; Institute of Clinical Research, University of Southern Denmark, Odense, Denmark; Wellcome Centre for Human Genetics, Nuffield Department of Medicine, University of Oxford, Oxford, United Kingdom; University of Witwatersrand, Johannesburg, South Africa; National Institute for Communicable Diseases, Johannesburg, South Africa; Department of Medical Microbiology, Centre for Infection and Immunity Amsterdam (CINIMA), Academic Medical Centre, University of Amsterdam, Amsterdam, The Netherlands; Netherlands Reference Laboratory for Bacterial Meningitis, Academic Medical Centre/RIVM, University of Amsterdam, Amsterdam, The Netherlands

## Abstract

*Streptococcus pneumoniae* is a common nasopharyngeal colonizer, but can also cause life-threatening invasive diseases such as empyema, bacteremia and meningitis. Genetic variation of host and pathogen is known to play a role in invasive pneumococcal disease, though to what extent is unknown. In a genome-wide association study of human and pathogen we show that human variation explains almost half of variation in susceptibility to pneumococcal meningitis and one-third of variation in severity, and identified variants in *CCDC33* associated with susceptibility. Pneumococcal variation explained a large amount of invasive potential, but serotype explained only half of this variation. Newly developed methods identified pneumococcal genes involved in invasiveness including *pspC* and *zmpD*, and allowed a human-bacteria interaction analysis, finding associations between pneumococcal lineage and *STK32C*.

---

*Streptococcus pneumoniae*, or the pneumococcus, is a leading cause of pneumonia, meningitis, and bacteremia. Over 90 serotypes are known, which have varying prevalence of asymptomatic carriage and disease^1,2^. Some clonal genotypes have been associated with invasive disease, though as serotype is correlated with genetic background, finding how much each of these factors affects invasive propensity in clinical cases of disease is challenging.

Bacterial meningitis involves severe inflammation of the membranes surrounding the brain, the meninges, which is a response to the presence of bacteria in the cerebrospinal fluid (CSF)^3^. *S. pneumoniae* is the most common cause of bacterial meningitis and despite advances in vaccination and treatment case fatality rate is 17-20% and unfavourable outcome occurs in 38-50% of cases^4^.

Knowledge of the contribution of genetic variability of humans and invading pathogens to pneumococcal meningitis susceptibility could guide development of new vaccines preventing the progression from asymptomatic carriage to invasive disease, whereas genetic variation associated with disease severity may guide new clinical intervention strategies during treatment^5^. However the effect of human genetics on pneumococcal meningitis is unknown – whether it affects the disease at all, and if so, which specific regions of the genome cause the effect. Historically, genetic association studies on bacterial meningitis have been held back by only assessing candidate genes, small sample sizes or poorly defined phenotypes^6^. More recent GWAS studies have found associations for children with meningococcal meningitis in Europe^7^, and pneumococcal meningitis in Kenya^8^.

In terms of pathogen variability, it is well known that the pneumococcal polysaccharide capsule, which determines its serotype, contributes to invasive propensity^1,2^. The pneumococcal genome also encodes a variety of proteins which directly interact with the host, mostly to enhance colonisation and avoid the host immune response^9^. Mouse models have shown that some antigens such *pspC* (*cbpA*) enhance virulence but are not essential in invasive isolates. Though the role of these antigens in colonisation and disease may be known, whether sequence variation at these loci has an effect on pathogenesis in human disease remains unclear. Previous small association studies have additionally suggested a role for platelet binding^10^ and arginine synthesis^11^ in pneumococcal meningitis, and analysis of within-host variation found that sequence variation of *dlt* and *pde1* are associated with pneumococcal meningitis^12,13^.

Thus, because of a lack of large cohort studies combined with detailed clinical metadata, the overall role of pneumococcal variation in clinical cases of meningitis is as yet unknown. It has not been possible to calculate the degree to which different serotypes affect invasiveness compared to other factors (either genetic or environmental), or if there are serotype-independent loci which are involved in invasion. Bacterial genome wide association studies (GWAS) provide a way to identify pneumococcal sequence variation associated with meningitis, independent of genetic background in an unbiased manner. While GWAS is more challenging in bacteria than humans due to strong population structure and high levels of pan-genomic variation, recent methodological advances have helped overcome these issues^14–16^.

We have collected data and samples from Dutch adults with meningitis between Jan 1, 2006, and July 1, 2014 in the prospective and nationwide MeninGene cohort^4^. Our study addresses previous limitations of genetic studies by collecting a large number of samples of both host and pathogen DNA from culture-proven cases of pneumococcal meningitis, along with detailed clinical metadata (supplementary table 1). We have performed genotyping and whole genome sequencing of this collection, a combined GWAS of host and pathogen in pneumococcal meningitis, the first time such a study has been attempted for a bacterial disease. We performed a GWAS separately in host and pathogen, continuing to develop new approaches to conduct the latter analysis. In both cases we have collected additional cohorts to replicate our findings (figure 1). We also had sequences available from both host and pathogen in 460 cases allowing analysis of interaction effects in a joint GWAS. Using these well characterised cohorts we explain the role of genetic variation of human and pathogen in pneumococcal meningitis.

**Table 1:** Human SNP heritability (*h*2SNP) of meningitis susceptibility and severity in Dutch adults. Heritabilities are shown on the liability scale (adjusted for population prevalence and case ascertainment ratio). We used two methods for each phenotype, GCTA and LDAK. The latter corrects for linkage disequilibrium when estimating the kinship between genotypes. All results showed significant evidence for a heritability above zero.

| Phenotype | Method | Heritability | Error | P-value |
| --- | --- | --- | --- | --- |
| Susceptibility (any species) | GCTA | 0.22 | 0.03 | $2.4 \times 10^{-6}$ |
| | LDAK | 0.29 | 0.05 | $3.9 \times 10^{-11}$ |
| Susceptibility (pneumococcal) | GCTA | 0.25 | 0.05 | $2.4 \times 10^{-6}$ |
| | LDAK | 0.29 | 0.07 | $3.9 \times 10^{-6}$ |
| Severity | GCTA | 0.29 | 0.11 | $2.8 \times 10^{-5}$ |
| | LDAK | 0.49 | 0.14 | $1.4 \times 10^{-4}$ |

**Figure 1:**
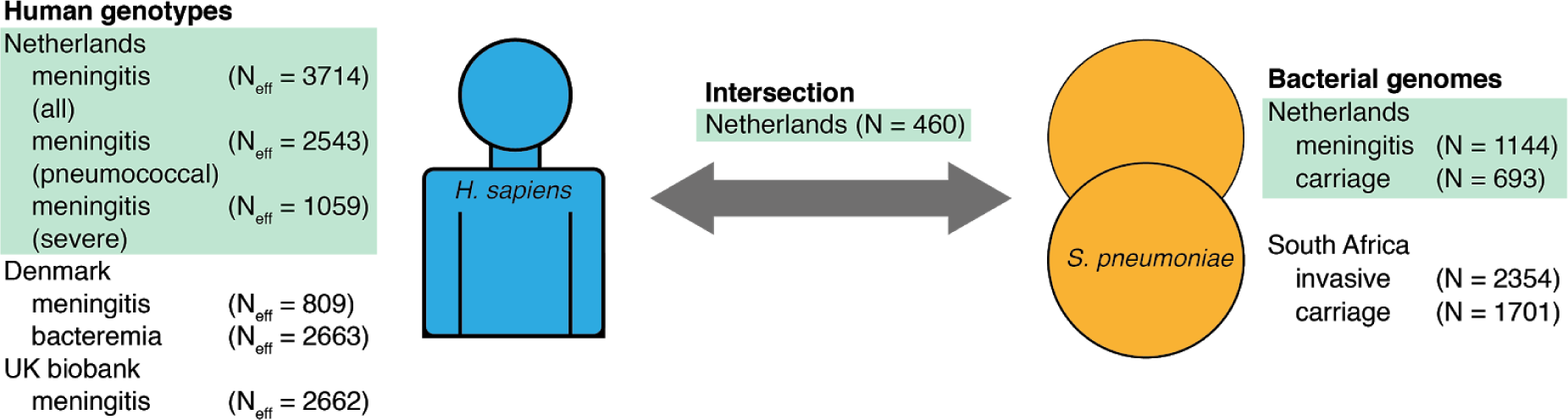
Overview of cohorts sequenced and associations performed. Left, host data; right, bacterial data; the centre represents samples with both host and pathogen sequence data. Samples in green are those collected from our MeninGene cohort that form the centre of this work. Due to unbalanced case control ratios we show the effective sample size, specific numbers of cases and controls of human genotypes are shown in Supplementary table 2.

## Results

### Human genetics influences pneumococcal meningitis susceptibility and severity

Before trying to find specific loci associated with meningitis, we first calculated the heritability of susceptibility and severity due to host genetics using the MeninGene cohort. While genetic associations have been found with invasive meningococcal disease^7^ and susceptibility to infectious diseases more broadly^17,18^, the heritability of adult meningitis and its outcome have not previously been calculated. We used two methods to calculate SNP heritability, which showed that variation in host genetics explains 29% ± 7% of the observed variation in pneumococcal meningitis susceptibility, and 49% ± 14% of the variation in meningitis severity (table 1).

**Table 2:** Signals of human association in the meningene cohort. we report the lead snp at each associated locus with maf > 5% and p < 1×10^−6^, and nearby annotated genes. the suggestive signal in all meningitis cases was also present when restricted to pneumococcal cases, albeit with a lower p-value of 3.9×10^−7^.

| Position (SNP) | Effect allele | MAF | OR | P-value | Annotation |
| --- | --- | --- | --- | --- | --- |
| **Susceptibility (any species)** |  |  |  |  |  |
| chr6:153582990 (rs3870369) | T | 0.42 | 1.27 | $7.2 \times 10^{-8}$ | Upstream of *RGS17* |
| **Susceptibility (pneumococcal)** |  |  |  |  |  |
| chr6:117624549 (rs210967) | G | 0.46 | 0.77 | $8.8 \times 10^{-7}$ | *ROS1* intronic |
| chr18:48403560 (rs2850542) | T | 0.43 | 0.65 | $7.6 \times 10^{-8}$ | *ME2* promoter (2kb upstream of TSS) |
| chr22:47506160 (rs13057743) | G | 0.33 | 0.74 | $5.5 \times 10^{-7}$ | *TBC1D22A* intronic |
| **Severity** |  |  |  |  |  |
| chr1:64680775 (rs12081070) | A | 0.43 | 1.62 | $2.0 \times 10^{-8}$ | *UBE2U* (5th intron)/*ROR1* |
| chr4:182823804 (rs2309554) | A | 0.33 | 1.58 | $4.1 \times 10^{-7}$ | *AC108142.1* intron |
| chr9:37382231 (rs72739603) | A | 0.07 | 2.36 | $6.7 \times 10^{-7}$ | *ZCCHC7/GRHPR* |

Finding that these traits were heritable we then used GWAS to search for specific loci associated with meningitis. Using the MeninGene cohort alone, one marker reached significance when testing for severity: position 64680775 (rs12081070) on chromosome 1, an intronic variant in *UBE2U* which was associated with unfavourable outcome (MAF = 0.43; OR = 1.63; p = 2.0×10^−8^) (supplementary figures 1-4). *UBE2U* is part of the ubiquitin pathway, but has not been previously associated with any disease or trait. Chromatin conformation capture data shows that the site of the most significantly associated variant interacts with *PGM1* and *ROR1* in a range of immune cell types including monocyte/macrophages, CD4/8 T cells, B cells and neutrophils (supplementary figure 5). *PGM1* encodes a phosphoglucomutase while *ROR1* is a protein of unknown function which has previously been associated with cancers^19^ and pulmonary function^20^. There was evidence of association of rs12081070 with gene expression in a panel of tissues and cell types but this was only significant in skin (p = 5.7×10^−13^)^21^. Six other loci showed suggestive significance (table 2), whereas in the Danish cohort, no variants reached genome wide significance (supplementary figures 6 & 7).

Of the genes implicated in the single cohort, we noted that the *ME2* promoter variant rs2850542 is an eQTL for the same gene in whole blood (p = 5.9×10^−20^, supplementary figure 8A) and specifically in monocytes (FDR 5.1×10-26)^22^. There was also evidence of chromatin interaction of the variant location with *SMAD4* again in a range of immune cell types including monocytes, lymphocytes and neutrophils (supplementary figure 8B/C). This gene is involved in TGF-β signalling and invasion across the epithelium; the variant also showed evidence of an eQTL involving rs2850542 for *SMAD4* expression in tibial artery (p = 8.6×10^−7^)^23^.

To improve our discovery power and mitigate false positives from batch effects, we then performed similar associations in the other cohorts. In an analysis of all available cases (MeninGene, Danish invasive disease, GenOSept, 23andme, UK biobank invasive disease) no hits were significantly associated with invasive disease (supplementary figure 9). However, the results for susceptibility to meningitis (MeninGene, Danish meningitis, UK biobank ICD-10 code for meningitis) found that position 74601544 on chromosome 15 (rs116264669) was associated with the minor allele increasing susceptibility in all three studies (p = 4.4×10^−8^; MAF = 3%) (supplementary figures 10 & 11). This intronic SNP is located in *CDCC33*, a gene that has no prior association with infectious disease. *CDCC33* is expressed in whole blood and the brain^21^ although there is no evidence of an eQTL involving this variant or SNPs in linkage disequilibrium with it. The disease-associated variant is located in a genomic region that interacts with the immunoglobulin superfamily containing leucine rich repeat 2 gene *ISLR2* on chromatin conformation capture analysis in macrophages (supplementary figure 12A) and CD8 T-cells (supplementary figure 12B); moreover a variant in complete linkage disequilibrium (rs80140040) with the meningitis susceptibility SNP shows evidence of eQTL with *ISLR2* in a number of tissues including brain (p = 0.02). *ISLR2* shows highest expression in the brain (neural tissues, supplementary figure 12C) and plays a role in the development of the nervous system^24^ but remains poorly characterised notably in humans. Severity data had not been recorded in other cohorts, and there were too few cases that had resulted in death to allow using this as a proxy for unfavourable outcome in a meta-analysis.

The results from the heritability analysis and GWAS suggest that susceptibility to meningitis is caused by many SNPs with individually low effect sizes (a polygenic trait), and that this cohort size is underpowered to detect these. To determine whether we expected the addition of more samples to uncover new associations with a larger effect size, we used Bayesian mixture models to fit the distribution of effect sizes in the MeninGene cohort. The maximum posterior suggested that 89% of the SNP heritability was caused by 29 large effect size SNPs (oligogenic), with the remaining heritability explained by small effect size SNPs (supplementary table 3). This was further supported by a dynamic Bayesian model incorporating variance in small and large effects, which found support for a mixture of many small effects plus a small number of large effects (maximum posterior ρ = 0.50 - proportion of variance explained by sparse terms) (supplementary table 4).

**Table 3:** Signals of bacterial association in the combined Dutch and South-African cohorts. Genes significant (Bonferroni corrected P < 0.05) in a pooled analysis of both cohorts with any of the association approaches, ordered by p-value. Odds-ratios are with respect to carriage samples. The genes in bold in the top half of the table are immunogenic, and are have previous evidence for association with virulence. For Tajima’s D the effect size is the difference between D values, and for k-mers and LoF burden tests it is the odds-ratio. For some p-values the calculation only allows an upper bound to be produced. The locus tag in the ATCC 700669 reference is listed, along with the common gene name if available.

| Gene ID | Gene name | Method | AF | OR/Effect size | P-value | Function |
| --- | --- | --- | --- | --- | --- | --- |
| SPN23F05680 | *ldcB/dacB* | Missense burden | 0.81 (gene) | 1.20 | $1.3 \times 10^{-19}$ | D-alanyl-D-alanine carboxypeptidase (peptoglycan peptide precursors) |
| SPN23F22240 | *pspC/cbpA* | K-mers | 0.2 (k-mer)<br>0.59 (gene) | 1.05 | $3.8 \times 10^{-13}$ | Binds secretory IgA, C3 and complement factor H; adhesin |
| SPN23F10590 | *zmpD* | Rare LoF in invasive | 0.53 (gene) | 1.42 | $<1 \times 10^{-10}$ | Unknown; paralogous to IgA1 protease (*zmpA*) |
| SPN23F08080 | *spnTVRhs dS* | Missense burden | 1.00 (gene) | 1.08 | $2.8 \times 10^{-6}$ | Type I restriction-modification system specificity subunit S |
| SPN23F17820 | ***psrP*** | Tajima's D | **0.42**<br>(gene) | **-2.71**<br>(invasive)<br>**-2.59**<br>(carriage) | **<1x10<sup>-6</sup>** | **adhesin** |
| SPN23F09820 |  | Missense burden | 0.36<br>(gene) | 1.3 | 3.4x10<sup>-49</sup> | Bacteriocin precursor |
| SPN23F12140 | *ntpl* | Missense burden | 0.88<br>(gene) | 0.71 | 1.3x10<sup>-48</sup> | V-type sodium ATP synthase subunit I |
| SPN23F05670 | *FM211187.1804* | Missense burden | 1.00<br>(gene) | 1.27 | 4.3x10<sup>-47</sup> | Histidine triad family protein (nucleotide phosphatase) |
| SPN23F04400 | *FM211187.1384* | Missense burden | 1.00<br>(gene) | 0.76 | 6.9x10<sup>-41</sup> | Unknown |
| SPN23F18990 | *FM211187.5748* | Missense burden | 1.00<br>(gene) | 0.76 | 1.9x10<sup>-18</sup> | ABC transporter ATP-binding protein |
| SPN23F01160 | *FM211187.339* | Missense burden | 1.00<br>(gene) | 0.88 | 2.7x10<sup>-11</sup> | Unknown |
| SPN23F20970 | *puuD* | Missense burden | 1.00<br>(gene) | 1.24 | 1.9x10<sup>-10</sup> | Gamma-glutamyl-gamma-aminobutyrate hydrolase |
| SPN23F04740 | *ecsA* | Missense burden | 1.00<br>(gene) | 1.15 | 1.4x10<sup>-8</sup> | ABC transporter ATPase |
| SPN23F16700 |  | Missense burden | 1.00<br>(gene) | 1.16 | 3.8x10<sup>-8</sup> | Nucleotide diphosphate hydrolase |
| SPN23F21080 | *phoR/pnpS* | Missense burden | 1.00<br>(gene) | 0.89 | 1.1x10<sup>-6</sup> | Phosphate-sensitive histidine sensor kinase |
| SPN23F11460 | *mcrB* | Missense burden | - | 1.13 | 1.2x10<sup>-6</sup> | Endonuclease |
| SPN23F05090 |  | Missense burden | 1.00<br>(gene) | 1.15 | 1.0x10<sup>-5</sup> | Aldose epimerase (putative) |
| SPN23F12730 |  | Missense burden | - | 0.77 | 2.1x10<sup>-5</sup> | Bacteriocin |

The collection of these datasets gave us the opportunity to perform two further multi-cohort analyses related to sepsis and self-reported meningitis. In the first, we combined Danish bacteremia, GenOSept and self-reported cases of sepsis in the UK biobank, but found no significant hits. Furthermore, we found no evidence that sepsis from clinician diagnosed cases in the UK biobank (using ICD-10 codes) were heritable. Secondly, we meta-analysed^23^ and me’s results for self-reported meningitis with self-reported cases in the UK biobank^17^ – the reported hit in *CA10* did not replicate. Given the p-value in the original study was just significant, it is possible that this result was a false positive. This may also be an artefact of respondent’s knowledge of whether they had bacterial meningitis, which requires expert knowledge to diagnose and distinguish from viral meningitis^25^.

### Multiple bacterial loci determine pneumococcal invasive potential

We first performed a heritability analysis to quantify the amount of variation due to the pneumococcal genome for each phenotype on the liability scale. We found that additive pneumococcal genetics explained much of variation in invasive propensity (*h*^2.^ = 85%), but no evidence of heritability of meningitis severity (*h*^2.^ = 0%). This suggests that invasive propensity is highly heritable, but that disease outcome is not determined by natural variation of pathogen genetics. The latter is not surprising as invasive disease is an evolutionary dead end for the pathogen so adaptations affecting virulence over the short course of infection are unlikely to be selected for. This is contrary to smaller studies which have suggested that bacterial genotype may help diagnose severe disease in a clinical setting^26^, but consistent with a meta-analysis finding no effect of pneumococcal serotype on the risk-ratio of death from meningitis^27^.

That pneumococcal invasiveness is affected by pneumococcal genetics is well known, but a quantitative estimate of by how much is unknown. The high heritability estimated here, suggests that in this population some bacteria are able to invade while others are not. This is consistent with some serotypes rarely found in invasive disease2, and with wide genetic separation from invasive lineages. The current focus of pneumococcal vaccination, and the most well known invasiveness determinant, is serotype. We therefore calculated what proportion of variation in invasiveness can be attributed to the serotype. Although not adjusted directly for genetic background, logistic regression gave a variance in invasiveness explained by the observed serotypes using Nagelkerke’s pseudo R2 from logistic regression of 0.45, less than the total heritability.

### Identification of pneumococcal invasiveness loci in multiple cohorts

Given that serotype does not explain all of the variation in invasive potential other pneumococcal factors associated with disease are likely to exist. We then looked at overall differences in the sequence variation between asymptomatic carriage and meningitis isolates. The amount of rare variation compared to common variation present in a population is informative of recent selection and population size changes^28^. An overall difference may therefore be informative of different selection on regions of the genome depending on the niche. In figure 2a we plot the site-frequency spectrum by niche and predicted consequence. Across the range of common minor allele frequencies (MAFs) in both niches the proportion of synonymous/nonsynonymous/intergenic/loss-of-function (LoF) mutations is as previously observed^29^; at low frequencies there is an excess of potentially damaging variants. Figure 2c shows the overall burden of damaging rare variants between carriage and invasive samples; in both LoF and damaging variants there was higher burden in carriage isolates (median LoF: invasive 7, carriage 11, W = 297440, p < 1×10^−10^; median damaging: invasive 22, carriage 26, W = 345370, p = 8×10^−4^),

**Figure 2:**
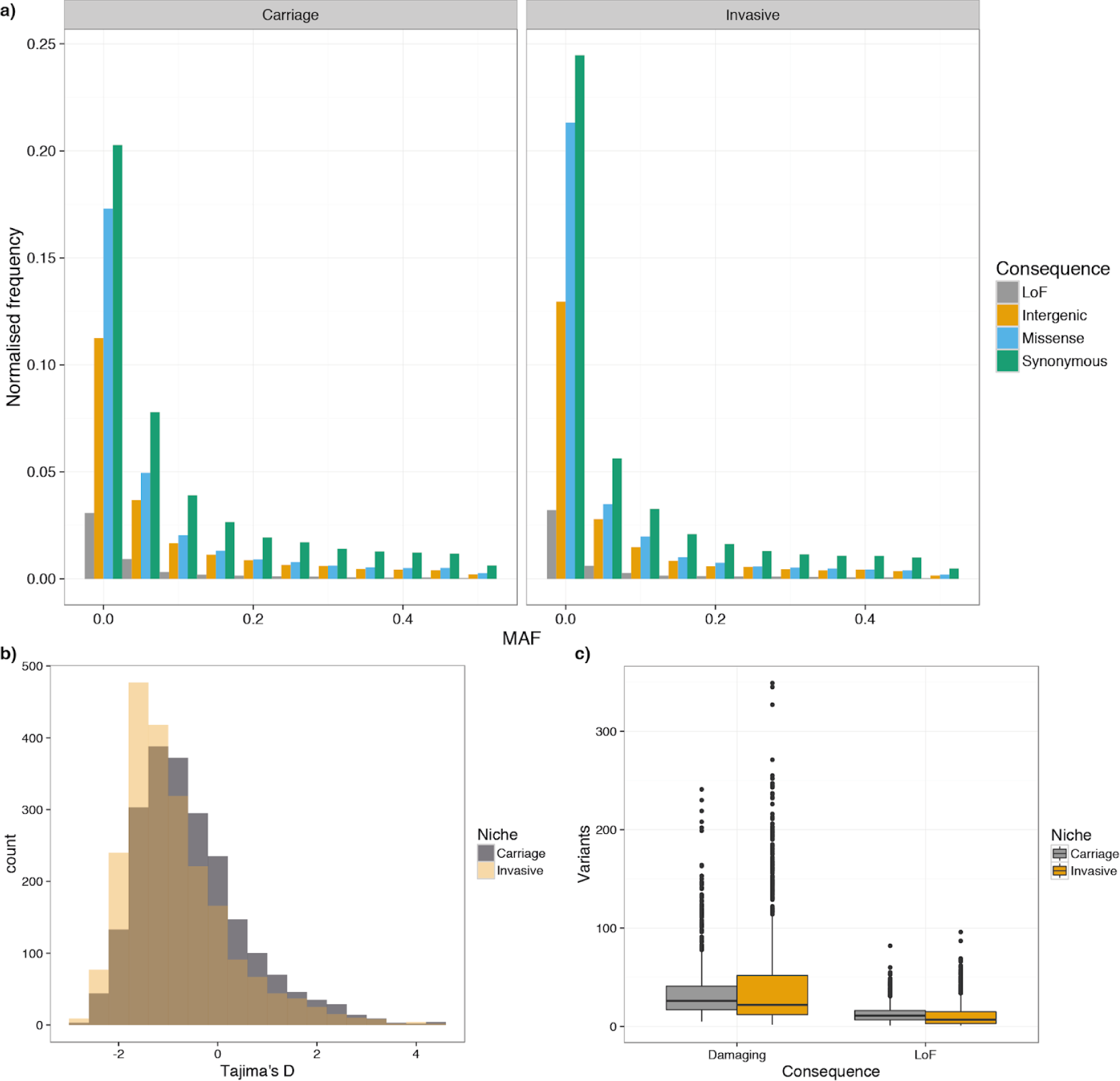
Differing burden and frequency of rare variation between invasive and carriage isolates, based on short variation called from mapping against the ATCC 700669 reference genome. Loss-of-function (LoF) are frameshift or nonsense mutations. a) The site frequency spectrum (SFS) stratified by niche and by predicted consequence. Frequency has been normalised with respect to the number of samples in each population. b) Histogram of Tajima’s D for all coding sequences in the genome, stratified by niche. c) Boxplot of number of rare variants per sample, stratified by niche and predicted consequence. Damaging mutations are LoF mutations and missense mutations predicted damaging by PROVEAN.

In total, invasive samples had a higher proportion of rare variants than carriage samples. To quantify this difference and identify which regions of the genome are responsible for the excess of rare alleles we calculated Tajima’s D, a statistic for neutral evolution, for each coding sequence in the genome, and looked for differing signs of selection between cases and controls. Deviations with D < 0 are indicative of selective sweeps and/or recent population expansion, whereas D > 0 is indicative of balancing selection and/or recent population contraction. In terms of differences between site-frequency spectrum (SFS), a negative D manifests as an excess of rare variants whereas a positive D manifests as a uniform distribution. We compared the distributions of D by gene in each phenotype (figure 2b). Genes in invasive isolates had a lower average D (difference in medians −0.34; W = 1996100, p < 10^−10^) and a more positively skewed D (difference in skewness 0.30; 95% bootstrapped CI 0.17-0.44). This difference in D may be representative of a genuine difference in selection of variants in genes between niche, or may be due to a difference in recent population dynamics, for example due to the bottlenecks for invasion and transmission.

We then used GWAS to find locus effects on invasive disease, independent of serotype. As pneumococcal genomes vary extensively in their pan-genome^30^, as well as having phenotypically important mosaic structural variants^31^ and antigens alleles^32^, we used a combination of methods to catalogue the population level variation then tested for associations while adjusting for population structure. We first performed this GWAS with disease severity, which was only measured in the Dutch cohort. Consistent with our estimates of no heritability, we found no loci of any type to be significantly associated with severity.

We then analysed meningitis versus carriage isolates. We first performed this analysis in the Dutch cohort (supplementary tables 5-7) revealing that many of our results involved rare variation, which has largely been ignored in previous bacterial GWAS studies^16^. This variation may be more important in a disease like meningitis where there is no pervasive selection for the phenotype. To improve the power and reliability of our results, we combined our Dutch cohort of meningitis and carriage samples with a cohort collected in South Africa, which included samples from carriage and cases of invasive pneumococcal disease. This gave a total of 5845 pneumococcal genomes to analyse. Table 3 shows the genes which were significant in this combined analysis using any of our association methods.

The genes noted in bold font in table 3 are all immunogenic^32^, and have all previously been associated with pneumococcal virulence in animal models^33–36^, but this is the first time an association has been shown in patients with invasive disease. Previous conclusions, drawn from protein binding to the Laminin receptor, have suggested that *pspC* (*cbpA*) is necessary for meningitis^37^. We predicted that both forms of *pspC* were absent in 13 meningitis isolates, though on closer inspection of the summary statistics from mapping and assembly these may also be an unresolved form of allele 8. Using our database of clinical data we also found that all of these patients had a severe ear, nose or throat infection, suggestive for direct spread of bacteria rather than crossing the blood-brain barrier. Three patients had clear bone destruction and/or pneumocephalus, which is proof for direct spread of the infection, and one patient had a skull defect. We further tested whether the two major forms of PspC were associated with meningitis specifically, as has previously been suggested^38^, but did not find either to be overrepresented, when accounting for population structure. *dacB* is involved in preserving cell wall shape, and has shown to attenuate virulence in a mouse model of lung infection. *zmpD* is homologous to IgA1 protease (*zmpA*)^39^, and while it is immunogenic^32^, its function is unknown – these results suggest a role in human cases of disease. The other genes in table 3 have not previously been directly associated with virulence or invasive disease in *S. pneumoniae*.

### Interactions between host and pathogen genomes

It is possible that different host genotypes have varying susceptibility to different lineages, or strains carrying certain alleles. To test this we used the 460 samples where we had collected both human genotype and pneumococcal genome sequence (figure 1). In the first instance, to retain hypothesis-free approach of GWAS, we performed a host-pathogen interaction analysis between every pair of common bacterial variants and genotyped host variants. While we were able to perform the 2×10^10^ associations required, no pairs of loci surpassed the large multiple testing burden required by this analysis - a power calculation showed that we would have 80% power for finding an effect with MAF of 25% and OR of 4 (supplementary figure 13) in the absence of population structure. Through this approach, we can therefore rule out large single interaction effects (with OR > 4 and MAF > 25%) between host and pathogen in cases of meningitis.

Given the difficulty of reaching significance for this large number of tests, we then went on to consider regions with strong prior evidence for being involved in host-pathogen interaction. *S. pneumoniae* has many virulence factors, some of which are known to interact with specific human proteins9. We were interested in the interactions where the pneumococcal protein contains sequence variation, ideally somewhat independent of genetic background. These regions have a higher power to be detected in an association analysis, and the higher rate of variation is potentially a sign of diversifying selection, which may mean the variation is more likely to be associated with specific interactions with the human immune system. We tested for an association between host genotype and the allele of three antigens selected for their variability and immunogenicity: PspC (CbpA), PspA, and ZmpA. For all of the antigen alleles with enough observations (supplementary table 8) we performed an association against all imputed human variants as above, and using a more accurate imputation of the *CFH* region due to its potential relevance in these interactions.

None of the bacterial antigen alleles were significantly correlated with variants in their human interacting-protein counterparts at the suggestive level (p < 10^−5^). However, there were two associations of a *cbpA* allele reaching genome-wide significance elsewhere in the genome. Supplementary figure 14 shows a locuszoom plot of each of these associations. The first is between *cbpA*-8 and position 148788006 on chromosome 6 (MAF = 0.08; OR = 9.20; p = 4.1×10^−9^). This is in *SASH1*, which has previously been found to have decreased expression during meningococcal meningitis (https://www.ebi.ac.uk/arrayexpress/experiments/E-GEOD-11755/). The second is between pspC-9 and position 98891272 on chromosome 13 (MAF = 0.16; OR = 6.30; p = 3.6×10^−8^), in *FARP1*, a gene not previously associated with infectious disease. Of note, we could find no published evidence of chromatin confirmation capture interaction or eQTL effects with either of these human variants.

We attempted to reduce the multiple testing burden by reducing the dimension of the pathogen genotype, which takes advantage of the extensive genome-wide correlation between variants. To give a straightforward biological interpretation we used hierarchical Bayesian clustering to define lineages, and tested whether pathogen genotypes, so defined, were associated with host genotype. We ran associations with lineages with at least 10% of samples in the subphenotype (supplementary table 9). The only result reaching genome-wide significance was an association between cluster eight (serotypes 9N/15B/19A, which have no overall association with invasive disease over carriage) and variants on chromosome 10 (supplementary figure 15). The lead variant (rs10870273) is at position 134046136 on chromosome 10 (MAF = 0.27; OR = 4.28; p = 1.2×10^−8^) located in an intron of STK32C, a serine/threonine kinase highly expressed in the brain. The high effect size estimated for the interaction is consistent with the power predicted in supplementary figure 13.

## Discussion

How genetics can affect susceptibility to and severity of pneumococcal meningitis has not been systematically investigated. We used multiple independent cohorts to perform human and pathogen GWAS to investigate the genetics of pneumococcal meningitis. We found no evidence for pathogen genetics affecting the severity of disease, whereas human genetics explained 49% ± 9% of this variation. In our Dutch cohort, variation near *UBE2U* and *ROR1* was significantly associated with severity. Our findings suggest that sequencing of the pathogen is likely to be uninformative for predicting disease progression, whereas further investigation of the host (and these genes in particular) may lead to greater insights into the mechanism behind severe cases of meningitis.

Host genetics explained 29% ± 7% of variation in susceptibility to meningitis, and a pooled GWAS analysis with our Danish cohort and the UK biobank found an association at *CDCC33*. This gene does not have a currently known function which is related to immunity, so this association may provide a new avenue with which to approach host studies; functional genomic data suggest a further possible mechanism for the susceptibility associated variant through interacting at a distance with and modulating expression of the brain expressed leucine-rich repeat and immunoglobulin (LIG) family protein gene *ISLR2*. By analysing the distribution of effect sizes, we found that meningitis susceptibility is likely to be affected by both a small number of large effect variants (oligogenic), and a large number of small effect variants (polygenic). We found no evidence of association between pneumococcal meningitis and *CFH*, which has previously been associated with meningococcal meningitis^7^, nor at any other candidate locus. We were also not able to report a previous finding in *CA10* from self-reported meningitis status, pointing to a need for careful clinical phenotyping needed in studies of hard to diagnose diseases such as bacterial meningitis.

Research of the role of pneumococcal variation in invasive potential in large epidemiological studies has mostly been focused on serotype variation. The lack of cohorts with whole genomes and invasive phenotypes has not allowed determination of other virulence factors in human disease – it is only with large cohorts of whole genome sequences that contributions to pneumococcal phenotypes can be systematically attributed to serotype or other genetic variation. With our large collection of genomes we were able to determine that the bacterial genome is crucial in determining invasive potential, with serotype likely to be the main factor (45% of variance explained). We went on to perform a combined analysis using 5892 pneumococcal genomes from two independent cohorts to find specific variation associated with invasive disease. We found five genes independent of genetic background and serotype to be associated with invasive disease. This showed a role for the virulence genes *cbpA* (*pspC*), *dacB* and *psrP* in human disease for the first time, and a new association with the loss of expression of the immunogenic protein *zmpD*.

Our cohort also allowed a joint analysis of bacterial and human sequencing data. Though the high dimension of data required more samples to find interaction effects of modest effect sizes, through biologically guided dimension reduction we were able to show evidence for possible enrichment of certain pathogen genotypes in certain host genotypes.

Pneumococcal meningitis, although a clinically important area of study due to its poor prognosis, is difficult to study. It is a relatively rare disease, challenging to diagnose in a timely manner, and requires a well set-up study to record the causal pathogen. Severe cases of meningitis are rare and may be difficult to record due to the need to follow up patients. Although there are no more current cohorts available to validate our results in, our findings suggest more cases should be collected to find further associations. Further replication using in vivo models will be necessary to confirm our results associating genes not previously known to be involved in pneumococcal meningitis.

Phenotype heterogeneity, here between causal organism, age of host infected and immunocompromised status may make replication of results with a large number of samples difficult. While we attempted to address this by looking for differences between sub-phenotypes, our sample size was likely too low to draw firm conclusions. It has been proposed that one reason why common variants associated with variation in infectious disease may be hard to find is that over human history they would have been selected against, and are therefore purged from the population^18,40^. Future genetic studies of meningitis should continue to collect samples with detailed clinical data, including the causal pathogen, to minimise phenotype heterogeneity.

## Methods

### Human genotyping and quality control

We performed genotyping using the Illumina Omni array, and called genotypes from normalised intensity data using optiCall^41^. For data taken from other platforms, we merged cases and controls only at sites in the intersection of the genotyping arrays used. We then performed basic quality control steps to first remove low quality samples, then low quality markers^42^. Samples with a heterozygosity rate three standard deviations away from the mean, or over 3% missing genotypes were removed. Markers with over 5% missing genotypes, significantly different (p < 10^−5^) call rate between cases and controls, MAF < 1% or out of Hardy-Weinberg equilibrium (p < 10^−5^) were removed. Using an LD-pruned set of markers, we estimated sample relatedness with KING^43^, and removed any duplicate samples. Using the same set of markers, we used eigenstrat to perform a PCA to check sample ancestry (supplementary figure 16)^44^. Samples closer than third-degree relation and samples of non-European ancestry (which we defined as PC1 < 0.07) were removed for heritability analysis, initial association attempts and analysis with Subtest. We manually inspected intensity plots for any associated markers using Evoker^45^, and removed any miscalled sites. Finally, we removed markers significantly associated with control batch (p < 5×10^−8^).

All markers were reported with respect to the reference allele and coordinates of GRCh^37^. We imputed markers using the HRC as a reference panel with the Sanger Imputation Server^46–48^. For the Danish samples, we instead used the Michigan imputation server due to the decreased number of markers available from merging two different genotyping arrays^49^. For greater accuracy, we reimputed the *CFH* region was imputed using impute2 with 1000 Genomes and GoNL as reference panels^50–52^. We removed resulting markers with MAF < 1%, HWE p < 10^−5^ or INFO scores < 0.7 leaving 6.8M markers for association testing and heritability estimation.

### Association of human variation

Throughout, we used a Glasgow Outcome Scale score^53^ of anything less than five (any long term disability or death) to define unfavourable outcome. We performed the association study using bolt-lmm^54,55^, using the LD-pruned set of genotyped markers to estimate the kinship matrix, and then calculating association statistics for all genotyped and imputed sites passing the above quality control thresholds. For the Dutch samples we included whether the patient was immunocompromised as a fixed effect (10% of cases), assuming that no control samples were immunocompromised (1% population prevalence^56,57^). To estimate heritability, we used two methods: GCTA-GREML^58^ (as implemented in bolt-lmm) and LDAK v5^59^. For both, we only used samples passing the stricter thresholds for ancestry and relatedness. Estimates of heritability were transformed from the observed scale to the liability scale using a population prevalence of meningitis of 1×10^−3^. When using Subtest to search for genetic difference between subphenotypes^60^, we used default settings, using the weights per marker calculated by LDAK to adjust for LD, and 1500 draws of 400 subsamples to generate null distributions of the test statistic.

We did not find evidence of overall differences between pneumococcal meningitis and other bacterial meningitis (pseudo-likelihood ratio (PLR) = 0.25; p = 0.75) or between severity and susceptibility (PLR = 0.14; p = 1.00). In the Danish cohort, there was no evidence of difference between meningitis and bacteremia (PLR = 311; p = 0.60). However this technique may rely on relatively highly associated SNPs, which were not found with this few samples. Susceptibility to any meningitis had a significantly higher heritability than its sub-phenotypes, which also have heritability above zero. This is more consistent with some difference in genetic architecture between the phenotypes.

To infer the distributions of effect sizes on meningitis, we ran bayesR^61^ and the Bayesian Sparse Linear Mixed Model (BSLMM), as implemented in gemma^62^. For both models, we used the genotyped sites (~630k) in the Dutch data using all meningitis cases as the phenotype. For bayesR we used version 2 of the software, using the ‘-shuffle’ option to increase computational efficiency. We ran the MCMC in each case with default settings: 5×105 iterations, discarding the first 2×10^5^ as burn-in and sampling every tenth iteration. We ran BSLMM using a probit link function for 1×10^6^ iterations, discarding the first 10^5^ as burn-in and sampling every tenth iteration. In both cases we report the mean value of each hyperparameter in the posterior.

To perform associations using the UK biobank we used bolt-lmm, following the recommended protocol for analysing the available genetic data^55^. We extracted case samples with self-reported meningitis or sepsis/septicaemia (data-field 20002), with a diagnosis of meningitis (data-field 41202 having a value of G01, G001, G002, G003, G008, A170, A390 or A321 at least once), and with a diagnosis of sepsis (A403, A409, A408 or A40 at least once). We randomly selected 3000 control samples from the remaining samples which had passed the UK biobank genetic quality control, which allowed for quicker analysis with little impact on effective sample size. Using this sample, we removed genotyped markers with MAF < 0.001 or a missing rate > 0.1, and used this to estimate kinships in bolt-lmm. We used bolt-lmm to perform association analysis of every imputed SNP site, including participant age as a fixed effect.

We used METAL to perform meta-analysis between different sets of studies^63^. We used the effective sample size to weight the beta and SE from each set of summary statistics, also adjusting the beta values and standard errors produced by bolt-lmm (supplementary table 10). We only retained markers which had been successfully imputed in all studies, to avoid effects of varying sample size at each locus.

Chromatin conformation capture data was tested and presented using the Capture HiC Plotter^64^ and eQTL data from the GTEx Consortium^21^.

### Catalogue of bacterial variation

From the whole genome sequence data of bacteria in the cohort we called SNPs and short INDELs with respect to the ATCC 700669 reference^65^. We mapped reads with bwa mem^66^, marked optical duplicates with Picard, and called variants with GATK HaplotypeCaller^67^. For INDELs we used the recommended hard filters. For SNPs we used the recommended hard filters to create an initial call set. We then applied GATK VariantRecalibrator using the following call sites as true positive priors: the intersection of SNPs called by both GATK and bcftools (Q10; 90% confidence); filtered SNPs from a carriage cohort of Karen infants^68^ (Q5; 68% confidence); filtered SNPs from a carriage cohort of children in Massachusetts^30^ (Q5; 68% confidence). After quality score recalibration we used 99.9% recall as a cutoff for SNPs to maximise sensitivity, and annotated the predicted effect of all coding variation using the variant effect predictor^69^. We defined LoF variants as either stop gained or frameshift mutations. We used PROVEAN with a score cutoff of < −2.5 to predict whether non-synonymous SNPs affected protein function^70^.

We produced a core gene alignment using roary^71^, where we used reciprocal best BLAST^72^ to choose a cut-off of 95% BLAST ID that maximised the balanced accuracy. To produce a presence/absence matrix of accessory elements, to be used as variants in association testing, we mapped the annotated genes in each isolate to a manually curated reference set^32,73^. As a first pass we used cd-hit^74^, and then used blastp against the genes which were unclustered after the first pass.

We counted variable length k-mers with a minor allele count of at least ten using fsm-lite^14^. In the Dutch data there were 11.7M informative k-mers, with 2.6M unique patterns. Using the mapping to the ATCC 700669 reference, we used cn.mops^75^ to call CNVs. We used windows of 1kb, and used windows with support for a CNV in at least two samples. The number of reads mapping to each *ivr* allele in each sample was determined by using read pair information, as previously^12,31^.

While the k-mer approach should directly assay or tag most variation at the population level, the allelic variation of the pneumococcal antigens may not be well captured. For example, *pspC* can be difficult to assemble due to repeats and copy number variation^76^, and k-mers from *pspA* and *zmpA* may not map to each allele specifically due to orthologous and paralogous genes^39,77^. As these antigens have been shown to interact with the host immune system, we developed a way to classify the allele present in each isolate combining assembly and mapping statistics. For each antigen in each sample we mapped reads to a reference panel using srst2^78^, and aligned annotated genes from the assemblies using blastp. We used coverage, number of SNP mismatches, number of INDEL mismatches and number of truncated bases as summary statistics for each reference allele from srst2, and equivalently percent ID, number of mismatches, number of gaps, E-value and bitscore from blastp. For *pspC*, we used an existing classification scheme of 11 alleles from 48 sequences^76^ (supplementary figure 17). For *zmpA* and *pspA* we built trees from previously characterised alleles^32^ (supplementary figures 18 & 19). Finding that the ancestral branches in the phylogenies were poorly supported, while the topology of clades were well resolved, we took a cut through the deep branches of the phylogeny to define four allele groups for *pspA* and three for *zmpA*. We ensured the training data were separable into these groups using PCA (supplementary figure 20). We tested the performance of four out-of-the-box classifiers on 20 *pspC* alleles spread across the population that we manually typed from the assemblies (supplementary table 11). Finding that an SVM with a linear kernel worked best, we used this to classify the allele of all antigens in all isolates using the summary statistics described above (supplementary figure 21).

Using the South African samples we counted k-mers, SNPs and INDELs and COGs in the same way, and annotated LoF function variants. We found 52215 SNPs and INDELs, 6.3M informative k-mers, with 1.5M unique patterns.

### Association of bacterial variation

Using the SNP and INDEL alignment we built a phylogenetic tree from this alignment using fasttree^79^, and calculated the kinship between each pair of strains as the distance between their MRCA and the midpoint root. We calculated heritability on the liability scale^80^ using this kinship matrix with FaST-LMM^81^. To estimate contributions for specific variant components such as serotype, we used lasso regression with leave-one-out cross validation to select significant predictors^16^ and Nagelkerke’s pseudo *R*^2^ to find the variance explained in phenotype^82,83^. To test the effect of capsule charge on phenotype we used previously measured zeta potentials in place of serotype^84^, using the serogroup average when a serotype specific charge was not known. Invasiveness was not well predicted from capsule charge alone (R2 = 0.08)^84^, partly because of the unknown capsule charge for many of the serotypes observed.

For association of common variation (MAF > 1%) we compared SEER^14^, using the first ten multidimensional scaling components as fixed effects to control for population structure, with FaST-LMM^81^, which uses eigenvectors from the kinship matrix calculated from the SNP and INDEL alignment as random effects to control for population structure. The Q-Q plots using fixed effects were highly inflated (supplementary figure 22), so we used the linear mixed model throughout. To correct for multiple testing we used the number of unique patterns as the number of tests in a Bonferroni correction, giving p < 8.2×10^−7^ for SNPs and p < 1.9×10^−8^ for k-mers. Inspection of the QQ-plots showed inflation of the test statistic for k-mers, so we used a higher threshold of p < 10^−16^ instead. The same association methods was used with the antigen alleles and CNVs.

We considered whether the *ivr* locus, a phase variable inverting type I R-M system with six possible alleles ^31^, is associated with meningitis. Using mouse models, alleles at this locus have previously been shown to be selected for in carriage, whereas others are preferred in invasive disease^35,85^. The rapid variation of this locus allows simple associations independent of genetic background. To test for association of the *ivr* locus alleles with susceptibility and severity, we used a Bayesian hierarchical model we had previously developed to find differences in the proportion of alleles present in tissue types while accounting for heterogeneity within single colony picks^12^. We used the same priors (using allele prevalence calculated using long-range PCR from a subset of samples) and MCMC parameters as specified previously, but labelling with phenotype rather than tissue. We found no evidence that either allele frequencies or overall diversity had any association with invasive disease or carriage in clinical cases of meningitis (supplementary figures 23 & 24).

To calculate Tajima’s D between phenotype groups we wrote a program in C++ using a function to calculate pairwise differences between strains we had previously optimised. Gaps or unknown sites were ignored. We calculated *D* for all coding sequences annotated in the ATCC 700669 reference, and p-values for difference between niche were calculated using 44000 null permutations of phenotype labels^86^. We applied a Bonferroni correction using the number of coding sequences tested.

Rare variants (MAF < 1%) could not be directly associated. Instead we applied a burden test^87^, grouping variants by coding sequence. As burden tests lose power when variants have different directions of effect on the phenotype, we used only those variants predicted to cause a loss of function in one test, and those causing either loss of function (6825 variants) or predicted change in protein function in another (additional 26206 variants). These variants will have occurred on terminal (or close to terminal) branches and therefore population structure is less of an issue than for common variants. We used plink/seq to perform this association for each phenotype, applying a Bonferroni correction using the number of genes as the number of multiple tests.

Reasoning that power to detect individual variants may also be hampered by population structure as well as allele frequency, we also searched for a burden of missense variants of any frequency by gene. We used the LMM burden testing mode of pyseer^88^ which allowed us to correct for population structure with the same model as above.

When performing analysis with both the Dutch and South African samples we pooled the genetic data, and performed the same association analysis as for the Dutch data alone. Where possible (associations using the linear mixed model) we included country as a covariate. The South African cohort also included host gender, age, collection year and HIV status and PCV-use at time of sampling. We included these as additional covariates for these samples. We were not able to use PROVEAN with this data, so only performed a burden test of damaging LoF variants. We used a significance threshold of P < 0.05 in the pooled analysis, after applying a Bonferroni correction for multiple testing based on the number of unique patterns as above. For all tests we ensured that the QQ-plots of the resulting p-values were not inflated (supplementary figure 25). We observed hits to simple transposons without cargo genes, which we therefore discarded as we assumed this was an artefact of their independence from population structure. We also found significant association of some BOX repeats^89^, but as there are many copies we could not map these associations to a single region of the genome.

### Interaction effects

We took 460 pneumococcal meningitis samples with matched pathogen and human sequence data which passed quality control thresholds for both data types. This has previously been applied to coding changes and host genotypes for HIV^90^ and HCV^91^, though these have much less variation than the pneumococcal population. To test all variants in a pairwise manner we converted the VCFs of human and pathogen calls into CSV files treating human genotypes as additive, and storing site and sample data separately for more efficient access by chunk^92^. The number of pairwise tests between all common variants was prohibitively large (10^12^), so we only tested genotyped markers: 1.8×1010 pairs of variants passed filters of MAF > 5% and missing rate < 5% in both the human and pathogen data. We modified the association code of SEER ^14^ to extend the 2 test to a 3×2 table, and to perform a 3×2 Fisher’s exact test (using https://github.com/chrchang/stats) when assumptions of the χ^2^ test were violated. Those sites with p < 5×10^−11^ (a Bonferroni correction with alpha = 1, as an initial filter) were then tested using a logistic regression of the human SNP against the pathogen variant, with the first three components from multidimensional scaling of the pathogen kinship matrix included as covariates to adjust for pathogen population structure.

To test for an association between invading lineage and human genotype we re-ran hierBAPS on the 460 samples^93^, which generated ten top level clusters (including a bin cluster) seven of which were large enough to test (supplementary table 9). We used bolt-lmm as above, but used whether the invading genotype was a member of the BAPS cluster as cases. We tested association of antigen alleles with frequencies over 10% in the sampled population in the same way (supplementary table 8).

## Acknowledgements

We would like to thank Chao Tan and David Hinds from 23andme for sharing summary statistics from their association with self-reported bacterial meningitis. Thanks to Win Kit Man for performing DNA isolation of bacteria. We also thank all members of the GenOSept consortium for making their data available and Anna Rautanen in particular for her preparation of the genetic data. This study makes use of data generated by the Genome of the Netherlands Project (GoNL). Funding for the GoNL project was provided by the Netherlands Organization for Scientific Research under award number 184021007, dated July 9, 2009 and made available as a Rainbow Project of the Biobanking and Biomolecular Research Infrastructure Netherlands (BBMRI-NL). Samples where contributed by LifeLines (http://lifelines.nl/lifelines-research/general), The Leiden Longevity Study (http://www.healthy-ageing.nl;http://www.langleven.net), The Netherlands Twin Registry (NTR: http://www.tweelingenregister.org), The Rotterdam studies, (http://www.erasmus-epidemiology.nl/rotterdamstudy) and the Genetic Research in Isolated Populations program (http://www.epib.nl/research/geneticepi/research.html#gip). The sequencing was carried out in collaboration with the Beijing Institute for Genomics (BGI).

## Funding

Work at the Wellcome Trust Sanger Institute was supported by Wellcome Trust (098051). This work was also supported by grants from the European Research Council (ERC Starting Grant, proposal/contract 281156; https://erc.europa.eu/) to DvdB and the Netherlands Organization for Health Research and Development (ZonMw; NWO-Vidi grant 2010, proposal/contract 016.116.358 to DvdB; NWO-Veni Grant NWO-Veni grant 2012 proposal/contract 916.13.078 to MB; www.zonmw.nl/). The Netherlands Reference Laboratory for Bacterial Meningitis was supported by the National Institute for Health and Environmental Protection, Bilthoven (www.rivm.nl/). JAL was supported by a Medical Research Council studentship grant (1365620). NJC was supported by a Sir Henry Dale Fellowship, jointly funded by the Wellcome Trust and the Royal Society (grant number 104169/Z/14/Z). JNW is funded by grants from the United States Public Health Service (AI038446 and AI105168). The BPROOF cohort was funded by The Netherlands Organization for Health Research and Development (ZonMw, Grant 6130.0031), the Hague; unrestricted grant from NZO (Dutch Dairy Association), Zoetermeer; Orthica, Almere; NCHA (Netherlands Consortium Healthy Ageing) Leiden/Rotterdam; Ministry of Economic Affairs, Agriculture and Innovation (project KB-15-004-003), the Hague; Wageningen University, Wageningen; VUmc, Amsterdam; Erasmus Medical Center, Rotterdam; Unilever, Colworth, UK. Dutch Carriage studies at RIVM were funded by the Dutch Ministry of Health, Welfare and Sport. Genotyping for GOYA was funded by the Wellcome Trust (WT 084762MA). GOYA is a nested study within The Danish National Birth Cohort which was established with major funding from the Danish National Research Foundation. Additional support for this cohort has been obtained from the Pharmacy Foundation, the Egmont Foundation, The March of Dimes Birth Defects Foundation, the Augustinus Foundation, and the Health Foundation. Funding for the Danish Invasive Pneumococcal Disease Cohort was kindly provided by the Lundbeck Foundation, the Novo Nordisk Foundation, King Christian the 10th Foundation, Jacob Madsen’s Foundation, TrygVesta, Ebba Celinder’s Foundation, the Danish Medical Association Foundation, the Foundation for Advancement of Medical Science, the Augustinus Foundation, Peder Laurits Pedersen’s Foundation, A.P. Møller Foundation for the Advancement of Medical Science, the Danish Medical Research Council, Preben and Anna Simonsen’s Foundation, Ferdinand and Ellen Hindsgaul’s Foundation, the Hartmann’s Foundation and Dagmar Marshalls Fond. The GenOSept study was supported by the European Union (6th framework programme of RTD funding) and the National Institute for Health Research, through the Comprehensive Clinical Research Network. JCK was supported by NIHR Oxford Biomedical Research Centre and a Wellcome Trust Investigator Award (204969/Z/16/Z) and AJM was supported by a Wellcome Trust Fellowship with reference 106289/Z/14/Z. The funders had no role in study design, data collection and analysis, decision to publish, or preparation of the manuscript.

## Author contributions

Conceptualization: JAL, BF, PHCK, JP, AvG, AvdE, MCB, JCB, SDB, DvdB. Data curation: JAL, BF, PHCK, NEW, NJC, RAG, HJB, ZBH, LHÄ, AJM, TCM. Formal analysis: JAL, BF, PHCK, NEW, AHZ, AJM, JCK. Funding acquisition: JP, SM, TB, AvG, AvdE, MCB, JCB, SDB, DvdB. Investigation: JAL, BF, PHCK, NEW, MVS, AJM, TCM, JCK. Methodology: JAL, BF, PHCK, NEW. Project administration: AvdE, MCB, JCB, SDB, DvdB. Resources: MVS, HJB, NR, AJWM, EAMS, KT, ALW, LHvdB, WvR, JHV, ZBH, LFL, LCPGMdG, NMvS, LHÄ, TIAS, EAN, MdP. Software: JAL. Supervision: JNW, JP, SM, TB, AvG, AvdE, MCB, JCB, SDB, DvdB. Writing – original draft: JAL, SDB, DvdB. Writing – review & editing: all authors.

## Data and code availability

Code used for the analysis, along with phylogenies and predicted antigen alleles can be found at https://github.com/johnlees/meningene. Bacterial metadata, including ENA accession numbers, can be accessed at on figshare (doi:10.6084/m9.figshare.5915314).

## Conflict of interest statement

NJC and SDB were consultants for Antigen Discovery, Inc involved in the design of a proteome array for *S. pneumoniae*. EAMS reports grants from the pharmaceutical companies GlaxoSmithKline and Pfizer outside the submitted work. KT reports grants from Pfizer and consultancy fees from Pfizer paid to University Medical Centre Utrecht, both received outside the submitted work. DvdB received departmental honoraria for serving on a scientific advisory board for GlaxoSmithKline paid to the Academic Medical Center, outside the submitted work.

## Ethics statement

For the MeninGene study written informed consent was obtained from all patients or their legally authorized representatives. The studies were approved by the Medical Ethics Committee of the Academic Medical Center, Amsterdam, The Netherlands (approval number: NL43784.018.13). For bacterial carriage samples from the Netherlands, written informed consent was obtained from both parents of each child participant, and from all adult participants. Approvals for the 2009 and 2012/2013 studies in children and their parents (NL24116 and NL40288/NTR3614) and for the study in elderly adults (NTR3386) were received from a National Ethics Committee in the Netherlands (CCMO and METC Noord-Holland). For the 2010/2011 study a National Ethics Committee in the Netherlands (the STEG-METC, Almere) decided that approval was not necessary. The studies were conducted in accordance with the European Statements for Good Clinical Practice and the Declaration of Helsinki of the World Medical Association. The Danish Invasive Pneumococcal Disease Cohort was approved by Danish Data Protection Agency (record no. 2007-41-0229 and 01864 HVH-2012-046). Ethical permission was obtained from The Ethical Committee of The Capital Region of Denmark (H-B-2007-085 and H-1-2012-063). According to Danish Legislation, the Research Ethics Committee can grant an exemption from obtaining informed consent for research projects based on biological material under certain circumstances, and for this study such an exemption was granted.

## References

1. Brueggemann, A. B. et al. Clonal relationships between invasive and carriage Streptococcus pneumoniae and serotype- and clone-specific differences in invasive disease potential. J. Infect. Dis. 187, 1424–1432 (2003).

2. Hausdorff, W. P., Feikin, D. R. & Klugman, K. P. Epidemiological differences among pneumococcal serotypes. Lancet Infect. Dis. 5, 83–93 (2005).

3. Mook-Kanamori, B. B., Geldhoff, M., van der Poll, T. & van de Beek, D. Pathogenesis and pathophysiology of pneumococcal meningitis. Clin. Microbiol. Rev. 24, 557–591 (2011).

4. Bijlsma, M. W. et al. Community-acquired bacterial meningitis in adults in the Netherlands, 2006–14: a prospective cohort study. Lancet Infect. Dis. 16, 339–347 (2016/3).

5. Brouwer, M. C. et al. Nationwide implementation of adjunctive dexamethasone therapy for pneumococcal meningitis. Neurology 75, 1533–1539 (2010).

6. Brouwer, M. C. et al. Host genetic susceptibility to pneumococcal and meningococcal disease: a systematic review and meta-analysis. Lancet Infect. Dis. 9, 31–44 (2009).

7. Davila, S. et al. Genome-wide association study identifies variants in the CFH region associated with host susceptibility to meningococcal disease. Nat. Genet. 42, 772–776 (2010).

8. Rautanen, A. et al. Polymorphism in a lincRNA Associates with a Doubled Risk of Pneumococcal Bacteremia in Kenyan Children. Am. J. Hum. Genet. 2, 1092–1100 (2016).

9. Kadioglu, A., Weiser, J. N., Paton, J. C. & Andrew, P. W. The role of Streptococcus pneumoniae virulence factors in host respiratory colonization and disease. Nat. Rev. Microbiol. 6, 288–301 (2008).

10. Tunjungputri, R. N. et al. Phage-Derived Protein Induces Increased Platelet Activation and Is Associated with Mortality in Patients with Invasive Pneumococcal Disease. MBio 8, (2017).

11. Piet, J. R. et al. Streptococcus pneumoniae arginine synthesis genes promote growth and virulence in pneumococcal meningitis. J. Infect. Dis. 209, 1781–1791 (2014).

12. Lees, J. A. et al. Large scale genomic analysis shows no evidence for pathogen adaptation between the blood and cerebrospinal fluid niches during bacterial meningitis. Microb Genom 3, e000103 (2017).

13. Lees, J. A. et al. Within-Host Sampling of a Natural Population Shows Signs of Selection on Pde1 during Bacterial Meningitis. Infect. Immun. 85, (2017).

14. Lees, J. A. et al. Sequence element enrichment analysis to determine the genetic basis of bacterial phenotypes. Nat. Commun. 7, 12797 (2016).

15. Earle, S. G. et al. Identifying lineage effects when controlling for population structure improves power in bacterial association studies. Nature Microbiology 16041 (2016).

16. Lees, J. A. et al. Genome-wide identification of lineage and locus specific variation associated with pneumococcal carriage duration. Elife 6, (2017).

17. Tian, C. et al. Genome-wide association and HLA region fine-mapping studies identify susceptibility loci for multiple common infections. Nat. Commun. 8, 599 (2017).

18. Chapman, S. J. & Hill, A. V. S. Human genetic susceptibility to infectious disease. Nat. Rev. Genet. 13, 175–188 (2012).

19. Reddy, U. R., Phatak, S. & Pleasure, D. Human neural tissues express a truncated Ror1 receptor tyrosine kinase, lacking both extracellular and transmembrane domains. Oncogene 13, 1555–1559 (1996).

20. Lutz, S. M. et al. A genome-wide association study identifies risk loci for spirometric measures among smokers of European and African ancestry. BMC Genet. 16, 138 (2015).

21. GTEx Consortium et al. Genetic effects on gene expression across human tissues. Nature 550, 204–213 (2017).

22. Fairfax, B. P. et al. Innate immune activity conditions the effect of regulatory variants upon monocyte gene expression. Science 343, 1246949 (2014).

23. Clarke, T. B., Francella, N., Huegel, A. & Weiser, J. N. Invasive bacterial pathogens exploit TLR-mediated downregulation of tight junction components to facilitate translocation across the epithelium. Cell Host Microbe 9, 404–414 (2011).

24. Mandai, K., Reimert, D. V. & Ginty, D. D. Linx mediates interaxonal interactions and formation of the internal capsule. Neuron 83, 93–103 (2014).

25. Khatib, U., van de Beek, D., Lees, J. A. & Brouwer, M. C. Adults with suspected central nervous system infection: A prospective study of diagnostic accuracy. J. Infect. (2016). doi:10.1016/j.jinf.2016.09.007

26. Cremers, A. J. H. et al. The contribution of genetic variation of Streptococcus pneumoniae to the clinical manifestation of invasive pneumococcal disease. bioRxiv 169722 (2017). doi:10.1101/169722

27. Weinberger, D. M. et al. Association of serotype with risk of death due to pneumococcal pneumonia: a meta-analysis. Clin. Infect. Dis. 51, 692–699 (2010).

28. Yang, Z. Computational Molecular Evolution. (OUP Oxford, 2006).

29. Thorpe, H. A., Bayliss, S. C., Hurst, L. D. & Feil, E. J. Comparative Analyses of Selection Operating on Nontranslated Intergenic Regions of Diverse Bacterial Species. Genetics 206, 363–376 (2017).

30. Croucher, N. J. et al. Population genomics of post-vaccine changes in pneumococcal epidemiology. Nat. Genet. 45, 656–663 (2013).

31. Croucher, N. J. et al. Diversification of bacterial genome content through distinct mechanisms over different timescales. Nat. Commun. 5, 5471 (2014).

32. Croucher, N. J. et al. Diverse evolutionary patterns of pneumococcal antigens identified by pangenome-wide immunological screening. Proc. Natl. Acad. Sci. U. S. A. (2017). doi:10.1073/pnas.1613937114

33. Ogunniyi, A. D. et al. Contributions of pneumolysin, pneumococcal surface protein A (PspA), and PspC to pathogenicity of Streptococcus pneumoniae D39 in a mouse model. Infect. Immun. 75, 1843–1851 (2007).

34. Abdullah, M. R. et al. Structure of the pneumococcal l,d-carboxypeptidase DacB and pathophysiological effects of disabled cell wall hydrolases DacA and DacB. Mol. Microbiol. 93, 1183–1206 (2014).

35. Manso, A. S. et al. A random six-phase switch regulates pneumococcal virulence via global epigenetic changes. Nat. Commun. 5, 5055 (2014).

36. Shivshankar, P., Sanchez, C., Rose, L. F. & Orihuela, C. J. The Streptococcus pneumoniae adhesin PsrP binds to Keratin 10 on lung cells. Mol. Microbiol. 73, 663–679 (2009).

37. Orihuela, C. J. et al. Laminin receptor initiates bacterial contact with the blood brain barrier in experimental meningitis models. J. Clin. Invest. 119, 1638–1646 (2009).

38. van der Maten, E. et al. Streptococcus pneumoniae PspC Subgroup Prevalence in Invasive Disease and Differences in Contribution to Complement Evasion. Infect. Immun. 86, (2018).

39. Bek-Thomsen, M., Poulsen, K. & Kilian, M. Occurrence and evolution of the paralogous zinc metalloproteases IgA1 protease, ZmpB, ZmpC, and ZmpD in Streptococcus pneumoniae and related commensal species. MBio 3, (2012).

40. Casanova, J.-L. Severe infectious diseases of childhood as monogenic inborn errors of immunity. Proc. Natl. Acad. Sci. U. S. A. 112, E7128–37 (2015).

41. Shah, T. S. et al. optiCall: a robust genotype-calling algorithm for rare, low-frequency and common variants. Bioinformatics 28, 1598–1603 (2012).

42. Anderson, C. a. et al. Data quality control in genetic case-control association studies. Nat. Protoc. 5, 1564–1573 (2010).

43. Manichaikul, A. et al. Robust relationship inference in genome-wide association studies. Bioinformatics 26, 2867–2873 (2010).

44. Price, A. L. et al. Principal components analysis corrects for stratification in genome-wide association studies. Nat. Genet. 38, 904–909 (2006).

45. Morris, J. a., Randall, J. C., Maller, J. B. & Barrett, J. C. Evoker: a visualization tool for genotype intensity data. Bioinformatics 26, 1786–1787 (2010).

46. Loh, P.-R. et al. Reference-based phasing using the Haplotype Reference Consortium panel. Nat. Genet. 48, 1443–1448 (2016).

47. McCarthy, S. et al. A reference panel of 64,976 haplotypes for genotype imputation. Nat. Genet. (2016). doi:10.1038/ng.3643

48. Durbin, R. Efficient haplotype matching and storage using the positional Burrows-Wheeler transform (PBWT). Bioinformatics 30, 1266–1272 (2014).

49. Das, S. et al. Next-generation genotype imputation service and methods. Nat. Genet. 48, 1284–1287 (2016).

50. Howie, B., Marchini, J. & Stephens, M. Genotype imputation with thousands of genomes. G3 1, 457–470 (2011).

51. 1000 Genomes Project Consortium et al. An integrated map of genetic variation from 1,092 human genomes. Nature 491, 56–65 (2012).

52. The Genome of the Netherlands Consortium. Whole-genome sequence variation, population structure and demographic history of the Dutch population. Nat. Genet. 46, 818–825 (2014).

53. Jennett, B. & Bond, M. Assessment of outcome after severe brain damage. Lancet 1, 480–484 (1975).

54. Loh, P.-R. et al. Efficient Bayesian mixed-model analysis increases association power in large cohorts. Nat. Genet. 47, 284–290 (2015).

55. Loh, P.-R., Kichaev, G., Gazal, S., Schoech, A. P. & Price, A. L. Mixed model association for biobank-scale data sets. bioRxiv 194944 (2017). doi:10.1101/194944

56. van Veen, M. G. et al. National estimate of HIV prevalence in the Netherlands: comparison and applicability of different estimation tools. AIDS 25, 229–237 (2011).

57. Harpaz, R., Dahl, R. & Dooling, K. The Prevalence of Immunocompromised Adults: United States, 2013. Open Forum Infect Dis 3, (2016).

58. Yang, J., Lee, S. H., Goddard, M. E. & Visscher, P. M. GCTA: a tool for genome-wide complex trait analysis. Am. J. Hum. Genet. 88, 76–82 (2011).

59. Speed, D., Hemani, G., Johnson, M. R. & Balding, D. J. Improved heritability estimation from genome-wide SNPs. Am. J. Hum. Genet. 91, 1011–1021 (2012).

60. Liley, J., Todd, J. A. & Wallace, C. A method for identifying genetic heterogeneity within phenotypically defined disease subgroups. Nat. Genet. 49, 310–316 (2017).

61. Moser, G. et al. Simultaneous discovery, estimation and prediction analysis of complex traits using a bayesian mixture model. PLoS Genet. 11, e1004969 (2015).

62. Zhou, X., Carbonetto, P. & Stephens, M. Polygenic modeling with bayesian sparse linear mixed models. PLoS Genet. 9, e1003264 (2013).

63. Willer, C. J., Li, Y. & Abecasis, G. R. METAL: fast and efficient meta-analysis of genomewide association scans. Bioinformatics 26, 2190–2191 (2010).

64. Javierre, B. M. et al. Lineage-Specific Genome Architecture Links Enhancers and Non-coding Disease Variants to Target Gene Promoters. Cell 167, 1369–1384.e19 (2016).

65. Croucher, N. J. et al. Role of conjugative elements in the evolution of the multidrug-resistant pandemic clone Streptococcus pneumoniaeSpain23F ST81. J. Bacteriol. 191, 1480–1489 (2009).

66. Li, H. Aligning sequence reads, clone sequences and assembly contigs with BWA-MEM. 3 (2013).

67. Van der Auwera, G. A. et al. From FastQ data to high confidence variant calls: the Genome Analysis Toolkit best practices pipeline. Curr. Protoc. Bioinformatics 43, 11.10.1–33 (2013).

68. Chewapreecha, C. et al. Dense genomic sampling identifies highways of pneumococcal recombination. Nat. Genet. 46, 305–309 (2014).

69. McLaren, W. et al. Deriving the consequences of genomic variants with the Ensembl API and SNP Effect Predictor. Bioinformatics 26, 2069–2070 (2010).

70. Choi, Y., Sims, G. E., Murphy, S., Miller, J. R. & Chan, A. P. Predicting the functional effect of amino acid substitutions and indels. PLoS One 7, e46688 (2012).

71. Page, A. J. et al. Roary: rapid large-scale prokaryote pan genome analysis. Bioinformatics 31, btv421 (2015).

72. Ward, N. & Moreno-Hagelsieb, G. Quickly finding orthologs as reciprocal best hits with BLAT, LAST, and UBLAST: how much do we miss? PLoS One 9, e101850 (2014).

73. Corander, J. et al. Frequency-dependent selection in vaccine-associated pneumococcal population dynamics. Nature Ecology & Evolution 1 (2017).

74. Fu, L., Niu, B., Zhu, Z., Wu, S. & Li, W. CD-HIT: accelerated for clustering the next-generation sequencing data. Bioinformatics 28, 3150–3152 (2012).

75. Klambauer, G. et al. cn.MOPS: mixture of Poissons for discovering copy number variations in next-generation sequencing data with a low false discovery rate. Nucleic Acids Res. 40, e69 (2012).

76. Iannelli, F., Oggioni, M. R. & Pozzi, G. Allelic variation in the highly polymorphic locus pspC of Streptococcus pneumoniae. Gene 284, 63–71 (2002).

77. Hollingshead, S. K., Becker, R. & Briles, D. E. Diversity of PspA: Mosaic genes and evidence for past recombination in Streptococcus pneumoniae. Infect. Immun. 68, 5889–5900 (2000).

78. Inouye, M. et al. SRST2: Rapid genomic surveillance for public health and hospital microbiology labs. Genome Med. 6, 90 (2014).

79. Price, M. N., Dehal, P. S. & Arkin, A. P. Fasttree: Computing large minimum evolution trees with profiles instead of a distance matrix. Mol. Biol. Evol. 26, 1641–1650 (2009).

80. Lynch, M. & Walsh, B. Genetics and Analysis of Quantitative Traits. (Sinauer, 1998).

81. Lippert, C. et al. FaST linear mixed models for genome-wide association studies. Nat. Methods 8, 833–835 (2011).

82. International Schizophrenia Consortium et al. Common polygenic variation contributes to risk of schizophrenia and bipolar disorder. Nature 460, 748–752 (2009).

83. Hosmer, D. W., Jr., Lemeshow, S. & Sturdivant, R. X. Applied Logistic Regression. (John Wiley & Sons, 2013).

84. Li, Y., Weinberger, D. M., Thompson, C. M., Trzciński, K. & Lipsitch, M. Surface charge of Streptococcus pneumoniae predicts serotype distribution. Infect. Immun. 81, 4519–4524 (2013).

85. Li, J. et al. Epigenetic Switch Driven by DNA Inversions Dictates Phase Variation in Streptococcus pneumoniae. PLoS Pathog. 12, e1005762 (2016).

86. Winantea, J. et al. A summary statistic approach to sequence variation in noncoding regions of six schizophrenia-associated gene loci. Eur. J. Hum. Genet. 14, 1037–1043 (2006).

87. Li, B. & Leal, S. M. Methods for detecting associations with rare variants for common diseases: application to analysis of sequence data. Am. J. Hum. Genet. 83, 311–321 (2008).

88. Lees, J. A., Galardini, M., Bentley, S. D., Weiser, J. N. & Corander, J. pyseer: a comprehensive tool for microbial pangenome-wide association studies. Bioinformatics (2018). doi:10.1093/bioinformatics/bty539

89. Croucher, N. J., Vernikos, G. S., Parkhill, J. & Bentley, S. D. Identification, variation and transcription of pneumococcal repeat sequences. BMC Genomics 12, 120 (2011).

90. Bartha, I. et al. A genome-to-genome analysis of associations between human genetic variation, HIV-1 sequence diversity, and viral control. Elife 2, 1–16 (2013).

91. Azim Ansari, M. et al. Genome-to-genome analysis highlights the effect of the human innate and adaptive immune systems on the hepatitis C virus. Nat. Genet. (2017). doi:10.1038/ng.3835

92. Ganna, A. et al. Ultra-rare disruptive and damaging mutations influence educational attainment in the general population. Nat. Neurosci. 19, 1563–1565 (2016).

93. Cheng, L., Connor, T. R., Sirén, J., Aanensen, D. M. & Corander, J. Hierarchical and spatially explicit clustering of DNA sequences with BAPS software. Mol. Biol. Evol. 30, 1224–1228 (2013).

